# Excitatory cholecystokinin neurons in CA3 area regulate the navigation learning and neuroplasticity

**DOI:** 10.1101/2025.09.13.676045

**Authors:** Fengwen Huang, Abdul Baset, Stephen Temitayo Bello

## Abstract

Hippocampus, a key hub of neural circuits for spatial learning and memory, has attracted tremendous studies. Neuronal information processing in the hippocampus can be regulated by many types of neuropeptides. Cholecystokinin (CCK), the most abundant neuropeptide in the central nervous system which is involved in modulating neuronal functions, such as cognition, memory and neuroplasticity, is widely expressed in the hippocampus. However, whether local excitatory CCK neurons modulates hippocampal function is still unclear. In this study, we showed that CA1 pyramidal neurons receive projections from excitatory CCK neurons in area CA3 (CA3^CCK^ neurons). Subsequently, activation of the CA1-projecting CA3^CCK^ neurons triggers the release of CCK. Then, we found that activity of CA3^CCK^-CA1 neurons supports the hippocampal-dependent tasks. Furthermore, inhibition of CA3^CCK^-CA1 projections or knockdown of CA3^CCK^ gene expression markedly impaired the behavioral tasks and neuroplasticity. Taken together, these results may add to a better understanding of how neuromodulators regulate the neural functions in central nervous system.

## Introduction

Neuropeptides are expressed and secreted throughout the mammalian brain where they play key roles in modulating neuronal activities and behaviors (Swaab 1982; Merighi et al., 2011; De Wied 1997). Long-term potentiation (LTP) and long-term depression (LTD) have been considered cellular mechanisms correlated with learning and memory in the central nervous system (CNS) (Grasselli and Hansel 2014). Hippocampus is a unique brain region that is critically involved in learning and memory (Ranganath and Hsieh 2016), and how neuromodulations affect the function of the hippocampal system has attracted lots of interest (Gedankien et al., 2023; Broussard et al., 2016; Li and Gao 2016; Karayol et al., 2021).

During the past decades, many studies have identified the neuronal mechanisms that guide neuropeptides across the hippocampal function and examined whether neuropeptides undergo specific adaptations in response to external innervation. However, most investigations focus on the classical monoamine neuromodulators, including acetylcholine (Ach), dopamine (DP), norepinephrine (NE), serotonin (ST), etc (Gedankien et al., 2023; Broussard et al., 2016; Li and Gao 2016; Karayol et al., 2021). Ach can regulate neuronal excitability and synaptic transmission in the mammalian brain (Takács et al., 2018). Several studies have demonstrated that stress increases Ach release in a brain region-specific manner (Picciotto et al., 2012; Mineur et al., 2013). For example, stress-induced increase in Ach level in the rat hippocampus and cortical area. DP is a critical modulator in neuronal circuitry, which has been shown to be involved in a variety of behavioral phenomena in the hippocampus, such as episodic memory formation (Chowdhury et al., 2012), spatial learning (Kempadoo et al., 2016), and synaptic plasticity (Hamilton et al., 2010). DP innervation from the midbrain is mainly via the dopamine receptor (D1/D5) when it is released into the dorsal hippocampus (Cai and Ford 2018). Additionally, the role of NE in memory retrieval requires signaling through the β1-adrenergic receptor in the hippocampus (Chang et al., 2011). The hippocampus has one of the denser inputs of adrenergic terminals (containing NE) in the CNS, indicating that the adrenergic system plays a role in learning and memory (Goodman et al., 2021). Interestingly, high concentration of serotoninergic fibers in the forebrain is in stratum lacunosum-moleculare (SLM) of hippocampal areas CA1 and CA3, where the axons of layer III neurons in the entorhinal cortex form excitatory synapses with the distal apical dendrites of pyramidal cells (Cai et al., 2013). This temporoammonic pathway is required for some spatial recognition tasks and for long-term consolidation of spatial memory. All these studies implied that neuropeptides play a critical role in the hippocampal system and influence subsequent behaviors.

Cholecystokinin (CCK), one of the most abundant neuropeptides in the CNS (Ma and Giardino 2022), has not received much attention and its function in the hippocampal system has been generally underestimated. Although several reports have shown that CCK enhances the excitatory synaptic transmission in the hippocampus and improves learning performance (Wei et al., 2013; Reisi et al., 2015). However, the exact mechanism by which CCK regulates the hippocampus plasticity and hippocampus-dependent behaviors has not yet been fully elucidated. To address this knowledge gap, we adopted the transgenic mice, optogenetics, GPCR-based sensor, extracellular recording, chemogenetics, RNA interference technique, calcium recording and behavioral task to investigate: 1) the distribution profile of CCK-positive neurons in the CA3 area of dorsal hippocampus (DHP), 2) activation of CA3^CCK^ neurons secrets the CCK at hippocampal SC-CA1 synapses. 3) the causal relationship between the Ca^2+^ response of excitatory CA3^CCK^ neurons and hippocampal functions. This study should bring conceptual advances to the foundation of neuropeptide modulation and the studies of hippocampus-dependent learning and memory.

## Results

### Distribution profile of CCK-positive neurons in the DHP

Although many studies have well documented the role of inhibitory CCK in the hippocampus (Klausberger et al., 2005; Ali 2007; Hefft and Jonas 2005), the function of excitatory CCK in the hippocampus is still unclear. To address this issue, we developed the transgenic CCK Cre::Ai14 (Tdtomato) reporter mice that express the red fluorescent protein tdTomato upon Cre-mediated recombination. This mouse line is suited in examining the distribution profile of CCK positive neurons in the dorsal hippocampus (DHP) (**Figure 1A-B**). Interestingly, we found that the proportion of CCK positive neurons in area CA1 and CA3 of CCK Cre/Ai14 mice was around 8.50 % and 20.34 %, respectively (**Figure 1C-D**). We further examined the properties of these CCK neurons in areas CA3, which may send dense CCK^+^ projections to the area CA1. Interestingly, excitatory neuronal marker CamkIIα was intensively colocalized with Tdtomato expressing CCK neurons in the area CA3 (**Figure 1E**). Moreover. the CCK proteins also show high co-localization in excitatory neurons in CA3 area (**Figure 1F**).

**Figure 1.**
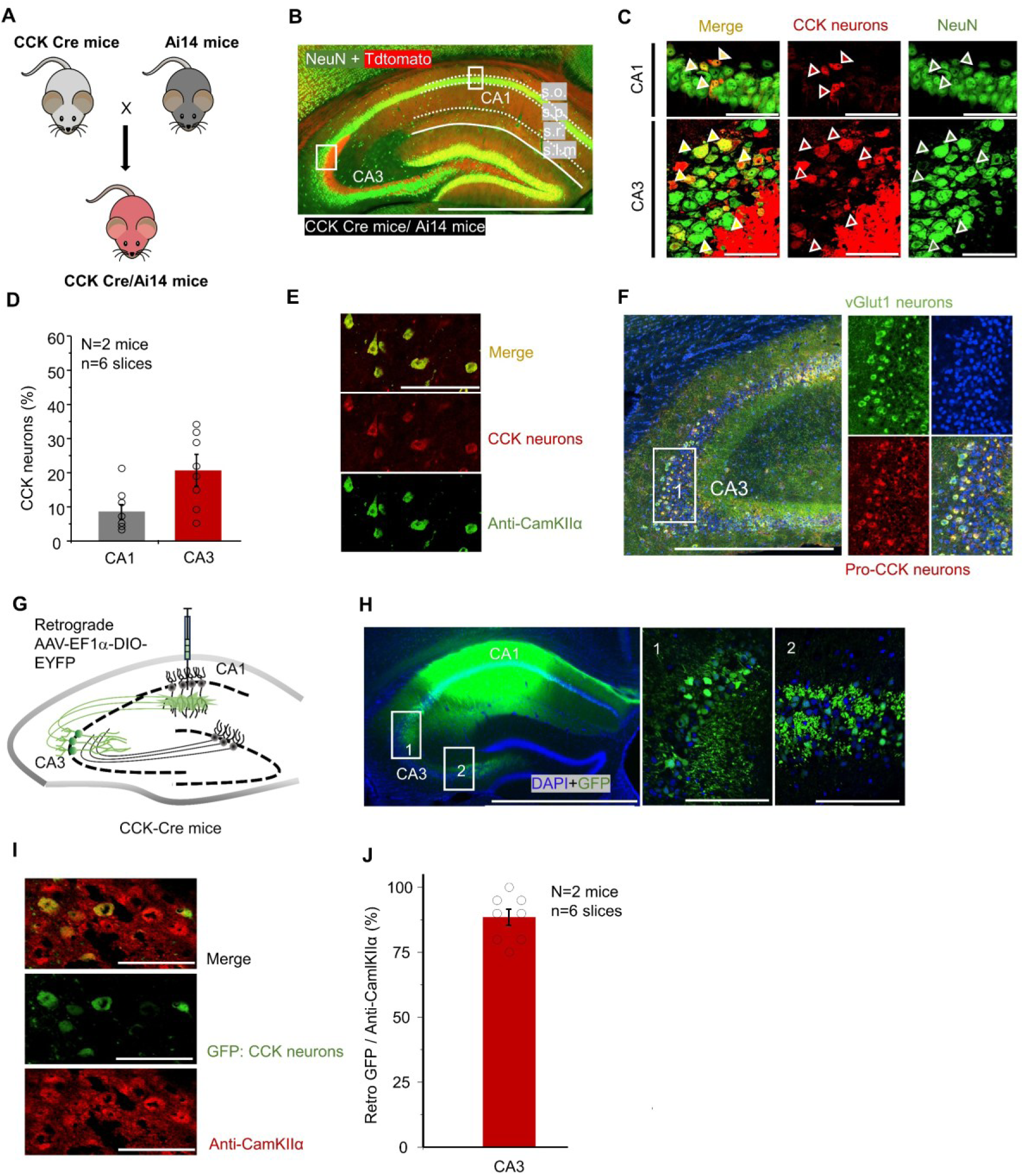
The distribution profile of CCK-positive neurons in the dorsal hippocampus. **(A)** Schematic illustrating the breeding strategy used to generate CCK-Cre/Ai14 reporter mice, in which CCK-expressing neurons are labeled with tdTomato. **(B)** Representative fluorescence image of a dorsal hippocampal slice from a CCK-Cre/Ai14 mouse immunostained with the pan-neuronal marker NeuN (rabbit anti-NeuN, Alexa Fluor 488), showing overall hippocampal anatomy. Scale bar, 1000 µm. **(C)** Higher-magnification images showing CCK-positive neurons identified by tdTomato expression (red) colocalized with NeuN immunofluorescence (green) in hippocampal CA1 and CA3 regions. Scale bar, 50 µm. **(D)** Quantification of the number of CCK-positive neurons in the CA1 and CA3 regions of the dorsal hippocampus. **(E)** Representative image of co-immunofluorescent labeling of CCK-positive neurons with the excitatory neuronal marker CaMKIIα in the CA3 region. Scale bar, 100 µm. **(F)** Fluorescence image showing colocalization of endogenous CCK protein (Pro-CCK) with an excitatory neuronal marker in the CA3 region. Scale bar, 1000 µm. **(G)** Schematic diagram illustrating the viral injection strategy. A Cre-dependent AAV (AAV-EF1α-DIO-EYFP; 6.5 × 10¹² vg/mL, 50 nL) was injected into the CA1 region of CCK-Cre mice to label CCK-expressing neurons and their projections. **(H)** Representative fluorescence images showing AAV expression in the hippocampal CA1 region (left) and retrogradely labeled neurons in the CA3 region (right). Scale bars: 1000 µm (left), 100 µm (right). **(I)** Representative image showing colocalization of retrogradely labeled GFP-positive neurons with the excitatory neuronal marker CaMKIIα (rabbit anti-CaMKIIα, red) in the CA3 region. Scale bar, 100 µm. **(J)** Co-immunofluorescent staining demonstrating colocalization of GFP with CaMKIIα in CA3 neurons of CCK-Cre mice, confirming the excitatory identity of retrogradely labeled CCK-expressing neurons. \**p□*<□0.05, \*\**p□*<□0.01, \*\*\**p□*<□0.001; ns not significant. Data are reported as mean□±□SEM.

Furthermore, to validate the projection of CA1-projecting CA3^CCK^ neurons in the DHP, we injected the Cre-dependent retrograde axonal transport of AAV (Retro-AAV-EF1α-DIO-EYFP) into the area CA1 (**Figure 1G**). Unsurprisingly, CA3 neurons were widely labelled with green fluorescent protein (GFP; **Figure 1H**). Moreover, high degree of colocalization between CamKIIα marker and GFP was observed in the area CA3 (**Figure 1I-J**; 90.15± 15 %). These anatomy results suggest that excitatory CA3 neurons are densely expressed and may exerts neuronal functions in central nervous system.

### Excitatory CA3 neurons can secret the neuropeptide CCK

Next, to investigate the neurotransmission property of the excitatory CCK neurons in CA3 area (CA3^CCK^), we utilized the Cre-dependent excitatory AAV (AAV9-DIO-CamKIIα-ChrimsonR-mCherry) to specifically target the excitatory CA3^CCK^ neurons in CCK-Cre mice (**Figure 2A**). The AAV shows high specificity for targeting the CCK positive neurons in the CA3 area (93.50 ± 1.97%; **sFigure 1**). Four weeks after the AAV injection and expression in *vivo*, robust expression of ChrimsonR-mCherry was observed in CA3^CCK^ neurons (**Figure 2B-C**). Subsequently, we used the 636 nm wavelength light to activate and elicit the field excitatory post synaptic potentials (L-fEPSPs) in ChrimsonR-mCherry expressing brain section (**Figure 2D**). Intriguingly, we noticed that the L-fEPSPs were almost completely blocked by the AMPA receptor antagonist (CNQX; 20 µM) and NMDA receptor antagonist (APV; 100 µM) in the area CA1, suggesting an excitatory nature of CA1-projecting CA3^CCK^ pathway (**Figure 2E-F**).

**Figure 2.**
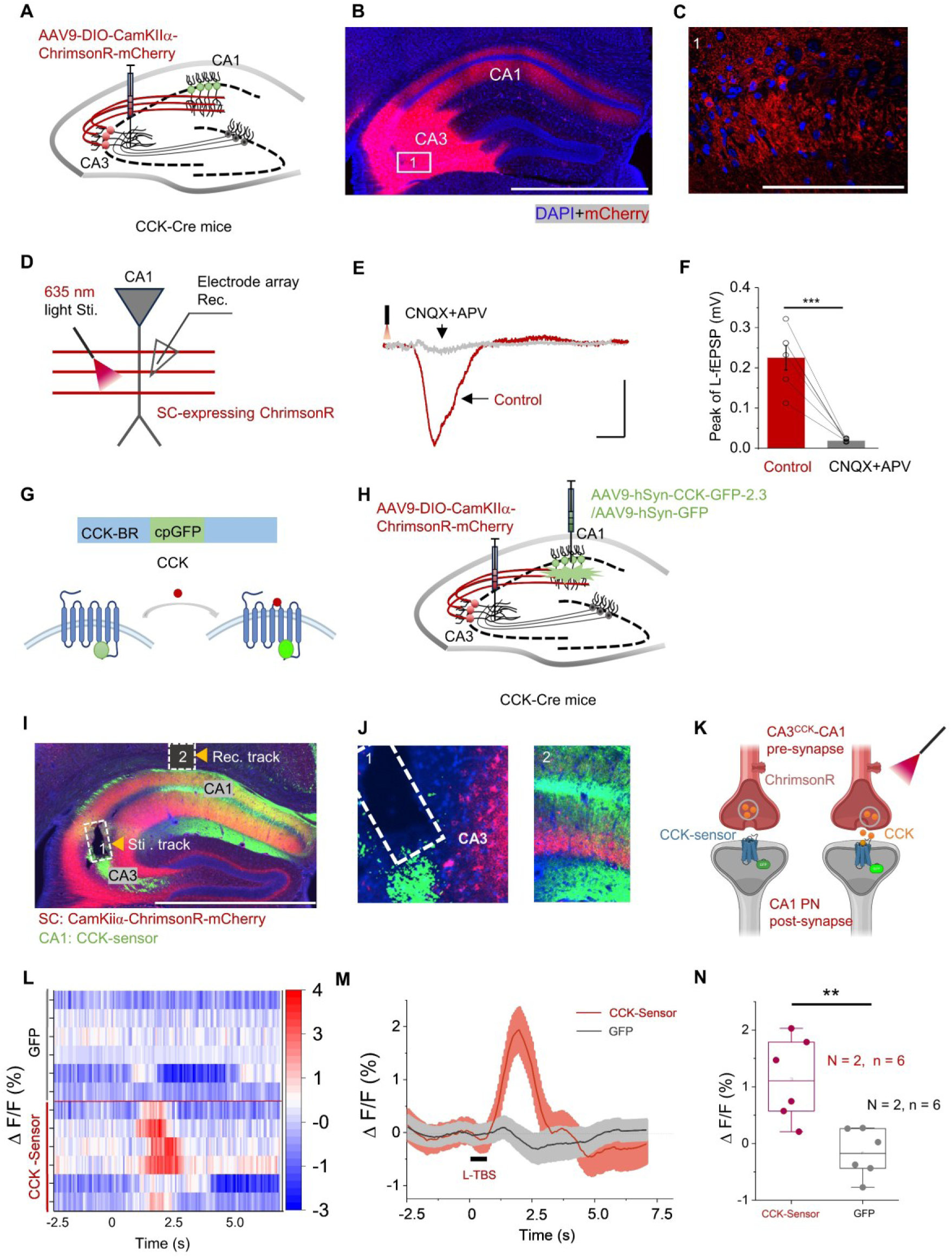
Excitatory CA3 neurons secret the neuropeptide CCK. **(A)** Schematic diagram illustrating the viral injection strategy. AAV9-DIO-CaMKIIα-mCherry (5.0 × 10¹² vg/mL, 250 nL) was injected into the CA3 region of CCK-Cre mice to selectively label excitatory CCK-expressing neurons. **(B)** Representative fluorescence image showing viral expression in the hippocampal CA3 region and labeled projections to the CA1 area. Scale bar, 1000 µm. **(C)** Higher-magnification image of the boxed region in (B), highlighting labeled CA3 axons projecting to CA1. Scale bar, 50 µm. **(D)** Schematic illustrating the experimental configuration for hippocampal slice electrophysiological recordings. **(E)** Representative light-evoked field excitatory postsynaptic potentials (L-fEPSPs) recorded from CA1 following optical stimulation of CA3 CCK-positive Schaffer collateral (SC) projections, shown before and after application of the glutamatergic receptor antagonists CNQX and APV. Scale bars, 0.2 mV and 10 ms. **(F)** Quantitative analysis of glutamatergic synaptic transmission mediated by CA3 CCK-positive projections to CA1, summarized across recordings. **(G)** Schematic illustration of the CCK sensor principle. Binding of the CCK ligand to the genetically encoded sensor induces a conformational change that results in altered fluorescence intensity. **(H)** Experimental schematic showing co-injection of AAV9-hSyn-CCK sensor 2.3 (5.75 × 10¹² vg/mL, 200 nL) and AAV9-CaMKIIα-DIO-ChrimsonR-mCherry (5.0 × 10¹² vg/mL, 250 nL) into the CA1 and CA3 regions, respectively, followed by optogenetic stimulation and fiber photometry recording. **(I)** Representative fluorescence image showing expression of the CCK sensor in the CA1 region surrounding the optical fiber tip (left; scale bar, 1000 µm) and ChrimsonR-expressing CA3-CA1 projections in CCK-Cre mice (right). **(J)** Higher-magnification images illustrating the optical stimulation site in CA3 and the photometry recording site in CA1. **(K)** Schematic model depicting activity-dependent release of CCK from CA3 presynaptic terminals onto CA1 neurons. **(L)** Heatmap representation of calcium-dependent fluorescence responses following optogenetic long theta-burst stimulation (L-TBS) in control GFP-expressing mice (upper; N = 3 animals, n = 6 trials) and CCK sensor-expressing mice (lower; N = 3 animals, n = 6 trials). **(M)** Averaged ΔF/F fluorescence responses evoked by optogenetic stimulation (635 nm L-TBS) in GFP and CCK sensor groups (N = 3 animals per group, n = 6 trials each). **(N)** Quantification of the mean fluorescence response in CCK-Cre mice, calculated as the averaged ΔF/F within 3 s following L-TBS. \**p□*<□0.05, \*\**p□*<□0.01, \*\*\**p□*<□0.001; ns not significant. Data are reported as mean□±□SEM.

Based on our recent work, CCK signaling in the hippocampus is predominantly mediated by CCK-B receptors, which play a critical role in regulating synaptic plasticity and spatial memory-related behaviors. To determine whether the neuropeptide CCK can be secreted from the CA3 derived excitatory CA3-CA1 projections, we adopted the GPCR-based CCK-BR sensor to monitor the release of CCK from the CA3-CA1 terminals in CA1 area under the external light stimulation (**Figure 2G**). To achieve this goal, AAV9-DIO-CamKIα-ChrimsonR-mCherry and AAV9-hSyn-CCK-GFP-2.3 (AAV9-hSyn-GFP as control) were injected into the CA3 and CA1 area to target the CA3-CA1 projection and CA1 pyramidal neurons (CA1 PNs) in CCK-Cre mice (**Figure 2H**). Subsequently, stimulation fiber and recording fiber were implanted upon the CA3 and CA1 area. Four weeks after the AAVs injection and expression (**Figure 2I-J**), high-frequency theta burst light stimulation (L-TBS) was delivered to the simulation fiber (**Figure 2K**), which mimic the neural rhythm in hippocampus in *vivo*. Interestingly, transient increase in CCK sensor was monitored by L-TBS of ChrimsonR-expressing CA3^CCK^ neurons, while no significant amplification was found in control (**Figure 2L-N**; Two samples T-Test, t = 3.79 df = 10, p = 0.004). These results indicates that neuropeptide CCK can be secreted from the excitatory CA3^CCK^ neurons under the mimic physiological conditions.

### CA3^CCK^ neurons fire actively during hippocampal-dependent tasks

We next wonder about the activity status of CA3^CCK^ neurons during hippocampal-dependent tasks (**Figure 3A)**. Thus, we injected the AAV9-CamkIIα-DIO-GCaMP6s into the area CA3 of CCK Cre mice to monitor the calcium-response of CA3^CCK^ neurons during the behavioral task (**Figure 3B**). The AAV displays high specificity for infecting the CCK positive neurons in the CA3 area (93.00 ± 1.33 %; **sFigure 2**). In novel object location (NOL) task, mice spent comparable time on object exploration between object 1 (O1) and object 2 (O2) in the training phase, while mice spent significantly more time on interacting with object in the novel place (O2”) compared with the object in the familiar site (**Figure 3C**; two way mixed ANOVA, Bonferroni adjustment; F_1,10_ = 4.58, p = 0.05; Training: O1 11.42 ± 1.56 s v.s. O2 10.62 ± 2.18 s, p = 0.73; Testing: O1 8.62 ± 0.99 s v.s. O2”: 13.95 ± 1.60 s, p = 0.02). Concurrently, we examined the Ca^2+^ responses of excitatory CA3^CCK^ neurons during object exploration. Interestingly, we observed that object exploration elicited a significant increase in fluorescence intensity in both training phase and testing phase (**Figure 3D-E**). Moreover, Ca^2+^signal of CA3^CCK^ neurons elicited by novel locations also show significant difference compared with familiar locations (**Figure 3F**; two way mixed ANOVA, Bonferrori adjustment; F_1,10_ = 1.88, p = 0.20; Training (Δ *F*/*F*): O1 0.27 ± 0.07 % v.s. O2 0.31 ± 0.08 %, p = 0.91; Testing (Δ *F*/*F*): O1 0.27 ± 0.09 % v.s. O2” 0.41 ± 0.12 %, p = 0.04), which implies that excitatory CA3^CCK^ neurons are critical for specific spatial memory and learning.

**Figure 3.**
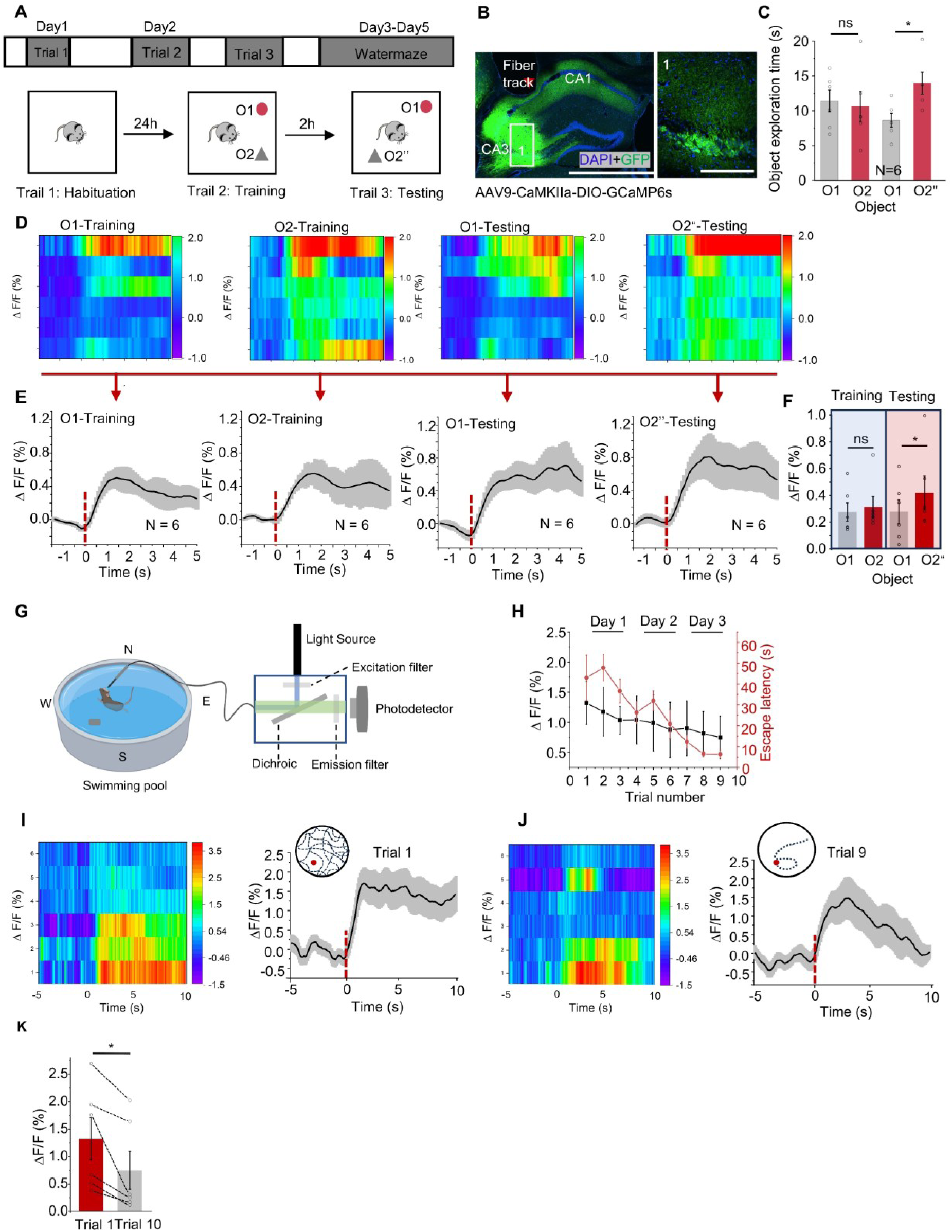
CA3^CCK^ neurons fire actively during hippocampal-dependent tasks. **(A)** Schematic overview of the behavioral paradigms used in this study, including the novel object location (NOL) task and the Morris water maze (MWM) task. **(B)** Representative fluorescence images showing viral injection into the CA3 region and expression of GCaMP6s in CCK-positive neurons of CCK-Cre mice following injection of AAV9-CaMKIIα-DIO-GCaMP6s (5.0 × 10¹² vg/mL, 300 nL). Scale bars, 1000 µm (left) and 100 µm (right). **(C)** Behavioral performance during the NOL task showing that mice spent significantly more time interacting with the object placed in a novel location compared with the familiar location. **(D)** Heatmap representation of ΔF/F fluorescence traces from a representative mouse during the training and testing phases of the NOL task (N = 6 mice). **(E)** Averaged ΔF/F fluorescence traces from all mice (N = 6), aligned to the onset of object exploration. **(F)** Mean GCaMP6s fluorescence signals of CA3 CCK-expressing neurons during object exploration bouts in training and testing trials, quantified as the average ΔF/F within 1.5 s following exploration onset. **(G)** Schematic illustration of the experimental setup for calcium imaging during the Morris water maze task. **(H)** Representative plot showing the relationship between normalized ΔF/F signals (black) and escape latency (red) across training trials in the MWM task. **(I)** Heatmap and mean GCaMP6s fluorescence signals during the initial learning phase (trial 1) of the MWM task. **(J)** Heatmap and mean GCaMP6s fluorescence signals during the well-trained phase (trial 9) of the MWM task. **(K)** Summary quantification of ΔF/F signals comparing trial 1 and trial 9, calculated as the average fluorescence within 10 s following trial onset. ^∗^p < 0.05, ^∗∗^p < 0.01, ^∗∗∗^p < 0.001; ns, not significant. Data are reported as mean ± SEM.

Furthermore, we examined whether the excitatory CA3^CCK^ neurons are necessary for performing the MWM paradigm (**Figure 3G**). Intriguingly, in the naive training (trial 1), a robust and persistent Ca^2+^ responses were observed during the initial learning (**Figure 3H and Figure 3I**). Nevertheless, after the completion of training, CA3^CCK^ neurons produced a relatively smaller Ca^2+^ signal on the training day 3 with an averaged escape latency of < 10 s to locate the hidden platform (**Figure 3H and Figure 3J**; **Figure 3K**: Paired sample t-test, df = 5, t = 2.80; Trial 1: 1.32 ± 0.74 % v.s. Trial 9: 0.75 ± 0.35 %, p = 0.038), this result further supports the conclusion that excitatory CA3^CCK^ neurons plays a direct role in the animals’ accurate spatial learning and memory formation.

### Chemogenetic inhibition of the excitatory CA3^CCK^-CA1 pathway impairs behavioral tasks

Next, to further confirm the necessity of excitatory CA3^CCK^-CA1 pathway for performance of the hippocampal-dependent behaviors. We adopted the chemogenetic approach, which is a powerful technique for specific disturbance of neuronal activity through virus-mediated DREADD expression in combination with its agonist clozapine N-oxide (CNO) (Gomez et al., 2017), to specifically target the excitatory CA3^CCK^ neurons. Firstly, we injected the Cre dependent AAV expressing the inhibitory DREADD (hM4DGi) to infect the excitatory CA3^CCK^ neurons and AAV carrying mCherry as control (**Figure 4A**) and implanted the drug cannular upon the CA1 area. The AAV exhibits high specificity for interacting the CCK positive neurons in the CA3 area (91.50 ± 2.24 %; **sFigure 3**). Four weeks after the AAV injection and expression (**Figure 4B**), we conducted the MWM task in the two groups of mice to assess the role of CA1-projecting CA3^CCK^ neurons in hippocampus-dependent spatial learning (**Figure 4C**). Thus, we delivered the CNO (300 nl; 10 µM) via the cannulas to inhibit CCK^+^ CA3-CA1 projections upon the area CA1 before conducting the MWM. Then, two groups of mice were subjected to the visible platform task to evaluate their swimming capability. Both groups of mice displayed comparable swimming speed and the speed time in finding the visible platform (**Figure 4D**; two sample t-test, df = 18, t = 0.22, HM4D(Gi): 24.3 ± 1.66 cm/s v.s. mCherry: 23.8 ± 1.52 cm/s, p = 0.83. **Figure 4E**; two sample t-test, df = 18, t = 1.09, HM4D(Gi): 30.86 ± 2.587 s v.s. mCherry: 34.21 ± 2.12 s, p = 0.29). During the hidden platform task, the mice only expressing mCherry gradually spent less time in locating the platform compared to the mice infected with hM4D(Gi) (**Figure 4F**; two-way mixed ANOVA, Bonferroni adjustment; F_4,15_ = 1.24, p = 0.34; HM4D(Gi): 24.18 ± 4.36 s v.s. mCherry: 15.12 ± 2.13 s on day 5, p = 0.003). Additionally, the control mice showed higher percentage on the target quadrant on the memory retention day, while lower percentage of the occupancy in experimental mice (**Figure 4G-H**; two-way mixed ANOVA, Bonferroni adjustment; F_3,16_ = 1.19, p = 0.34; HM4D(Gi): 27.91 ± 2.75 % v.s. mCherry: 42.31 ± 3.82 % in quadrant 3, p = 0.02). These results demonstrated that excitatory CA3^CCK^-CA1 pathway is required for animals to perform adequately in spatial learning. Additionally, high-frequency electrical stimulation fails to induce LTP in the CA3-CA1 pathway in both CCK-KO and CCK-BR-KO mice, indicating that CCK-dependent synaptic plasticity in this circuit is primarily mediated by CCK-B receptors.

**Figure 4.**
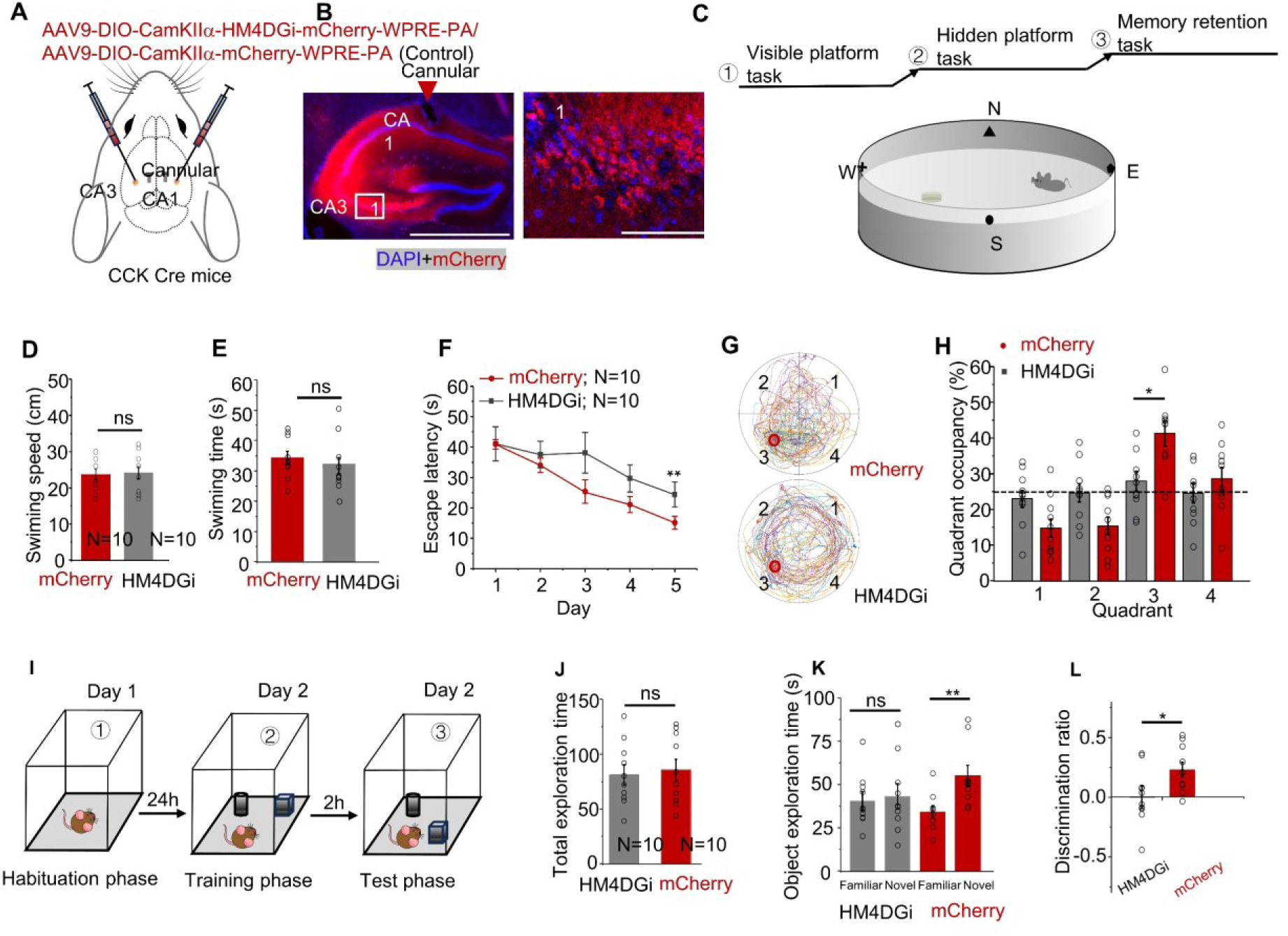
Chemogenetic inhibition of the excitatory CA3^CCK^-CA1 pathway impairs behavioral tasks. **(A**) Schematic illustrating Cre-dependent viral labeling and chemogenetic manipulation of CA3–CA1 projections in CCK-Cre mice. AAV9-CaMKIIα-DIO-hM4D(Gi)-mCherry (6.5 × 10¹² vg/mL, 300 nL) or control AAV9-hSyn-DIO-mCherry (6.5 × 10¹² vg/mL, 300 nL) was injected to selectively target excitatory CCK-expressing neurons. **(B)** Representative fluorescence image showing viral expression in the CA1 region with the corresponding cannula track. A higher-magnification image of the boxed area is shown on the right. Scale bars, 1000 µm (left) and 100 µm (right). **(C)** Schematic of the Morris water maze (MWM) task. The hidden platform was located in the southwestern (SW) quadrant (quadrant 3). **(D)** Quantification of swimming speed during the visible platform task, showing no significant differences between control and hM4D(Gi)-expressing mice. **(E)** Quantification of latency to locate the visible platform, indicating comparable sensorimotor performance between groups. **(F)** Escape latency across training days during the hidden platform phase of the MWM, comparing control and hM4D(Gi) groups. **(G)** Representative swimming trajectories of control and hM4D(Gi)-expressing mice during the spatial probe trial. **(H)** Percentage of total time spent in each quadrant during the probe test, showing reduced preference for the target quadrant in hM4D(Gi)-expressing mice. **(I)** Schematic of the novel object location (NOL) task (see Methods for details). **(J)** Total object exploration time during the NOL task, showing no significant difference between groups. **(K)** Control mice expressing mCherry exhibited a stronger preference for the object in the novel location compared with hM4D(Gi)-expressing mice. **(L)** Quantification of discrimination index demonstrating a significant reduction in spatial discrimination performance in hM4D(Gi)-expressing mice compared with controls. ∗p < 0.05, ∗∗p < 0.01, ∗∗∗p < 0.001; ns, not significant. Data are reported as mean ± SEM.

In the NOL task (**Figure 4I**), both groups of mice showed equal motivation to explore objects (**Figure 4J**; two sample t-test, df = 18, t = −0.31, hM4D(Gi): 81.18 ± 9.14 s v.s. mCherry: 85.35 ± 9.67 s, p = 0.76). Compared with the control mice, the experimental group was unable to distinguish the novel and familiar object location in the test phase, and presented low discrimination ratio (**Figure 4K**; two-way mixed ANOVA, Bonferroni adjustment; F_1,18_ = 4.58, p = 0.04; hM4D(Gi): Familiar 40.83 ± 5.27 s v.s. Novel: 43.46 ± 7.40 s, p = 0.66; mCherry 34.38 ± 3.58 s v.s. Novel: 55.35 ± 6.11 s, p = 0.003. **Figure 4L**; two sample t-test, df = 18, t = 0.77, hM4D(Gi): −0.003 ± 0.09 v.s. mCherry: 0.22 ± 0.06, p = 0.03), indicating suppression of excitatory CA3^CCK^-CA1 pathways impaired the spatial memory. Taken together, we can conclude that excitatory CA3^CCK^ neuron is an essential and functional component in the hippocampal system.

### Chemogenetic inhibition of the excitatory CA3^CCK^-CA1 pathway attenuates LTP formation

Since the LTP in the hippocampus is commonly considered a key cellular mechanism that underlies spatial learning and memory (Ge et al., 2010). To confirm whether excitatory CA3^CCK^-CA1 pathway are directly involved in LTP formation, we further conducted the electrophysiological recording in *vitro*. Four weeks after the AAV injection and expression (**Figure 5A-B**), hippocampal slices were subjected to electrophysiological recording to test the effect of CNO on the neuroplasticity of excitatory CA3^CCK^-CA1 pathway (**Figure 5C**).

**Figure 5.**
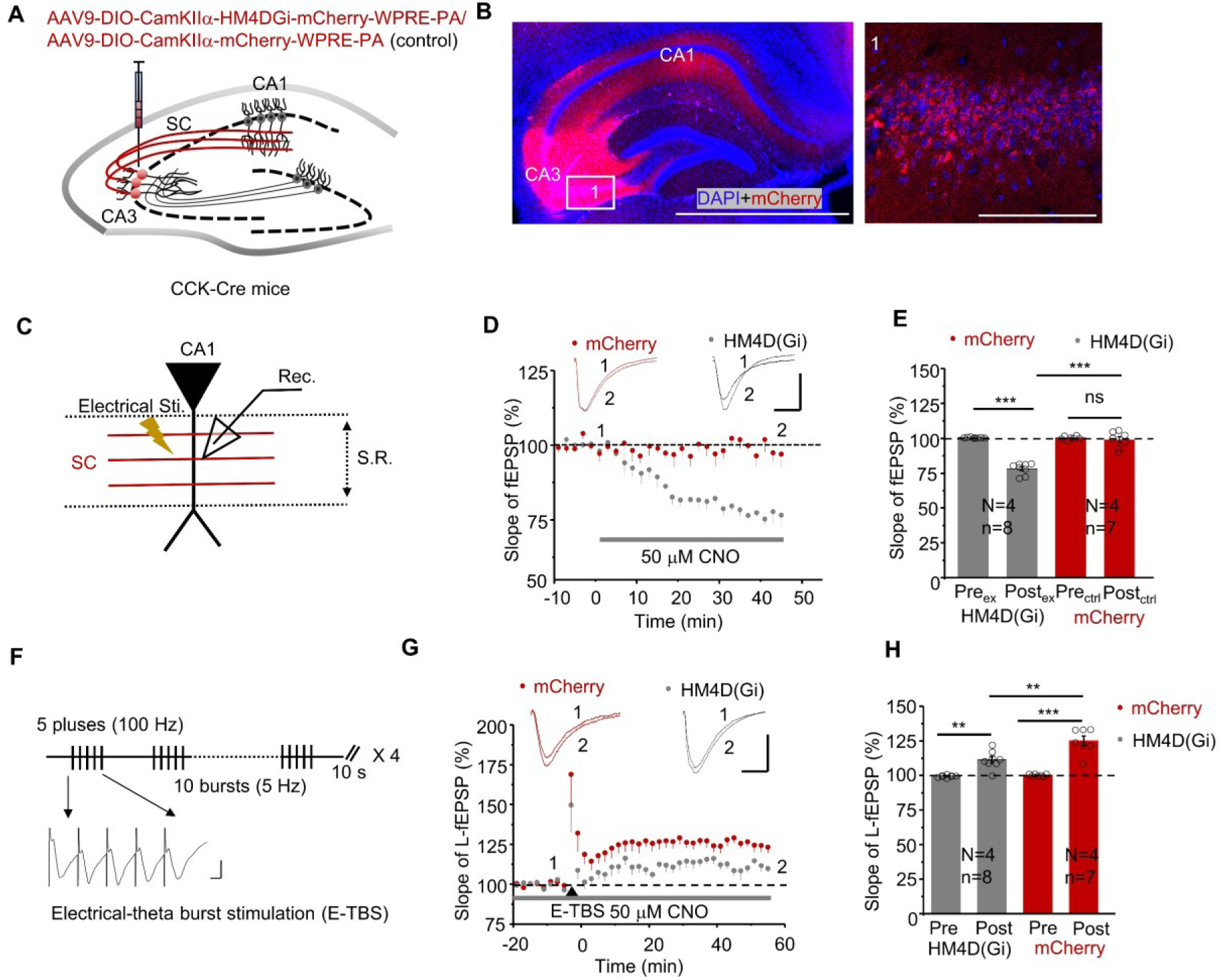
Chemogenetic inhibition of the excitatory CA3^CCK^-CA1 pathway impairs LTP formation. **(A)** Schematic illustrating the viral injection strategy. AAV9-CaMKIIα-DIO-hM4D(Gi)-mCherry (5.0 × 10¹² vg/mL, 300 nL) or control AAV9-hSyn-DIO-mCherry (5.0 × 10¹² vg/mL, 300 nL) was injected into the CA3 region of CCK-Cre mice to selectively target excitatory CCK-expressing neurons. **(B)** Representative fluorescence images showing viral expression in the hippocampal CA3 region and labeled projections to the CA1 area. A higher-magnification image of the boxed region is shown on the right. Scale bars, 1000 µm (left) and 100 µm (right). **(C)** Schematic illustration of the hippocampal slice electrophysiological recording configuration. **(D)** Representative electrically evoked field excitatory postsynaptic potentials (E-fEPSPs) recorded in CA1 following stimulation of CA3 Schaffer collateral inputs before and after application of clozapine-N-oxide (CNO). Chemogenetic inhibition significantly reduced E-fEPSP amplitude in slices expressing hM4D(Gi) compared with mCherry-expressing controls. Scale bars, 0.2 mV and 10 ms. **(E)** Quantification of E-fEPSP responses showing reduced synaptic transmission in hM4D(Gi)-expressing slices relative to control slices following CNO application. **(F)** Schematic of the electrical theta-burst stimulation (E-TBS) protocol used to induce long-term potentiation (LTP). **(G)** Representative traces showing that LTP induction was attenuated in hM4D(Gi)-expressing slices compared with mCherry controls. Scale bars, 0.2 mV and 10 ms. **(H)** Summary quantification of E-fEPSP responses following E-TBS, demonstrating significantly reduced LTP in the hM4D(Gi) group relative to controls. ∗p < 0.05, ∗∗p < 0.01, ∗∗∗p < 0.001; ns, not significant. Data are reported as mean ± SEM.

After the baseline recording, the slope of fEPSPs decreased gradually with the CNO treatment while no significant changes were observed in controls (**Figure 5D-E**; two-way mixed ANOVA, Bonferroni adjustment; F_1,13_ = 78.32, p < 0.001; hM4D(Gi): Pre_ex_ 100.42 ± 0.18 % v.s. Post_ex_ 78.07 ± 1. 52 %, p < 0.001; mCherry: Pre_ctrl_ 100.15 ± 0.48 % v.s. Post_ctrl_ 98.90 ± 2.90 %, p = 0.60, Post_ex_ v.s. Post_ctrl_, p < 0.001), indicating that neuronal activities of the excitatory CA1-projecting CA3^CCK^ neurons were remarkably suppressed. Then, we further employed the E-TBS protocol for the LTP induction under this condition (**Figure 5F**). Intriguingly, the extent of LTP formation in the slices expressing hM4D(Gi) was significantly smaller than those in the control group (**Figure 5G-H**; two way mixed ANOVA, Bonferrori adjustment; F_1,13_ = 9.46, p = 0.01; HM4D(Gi): Pre_ex_ 99.65 ± 0.31 % v.s. Post_ex_ 111.52 ± 2.40 %, p < 0.001, mCherry: Pre_ctrl_ 100.41 ± 0.29 % v.s. Post_ctrl_ 125.17 ± 3.43 %, p = 0.001, Post_ex_ v.s. Post_ctrl_, p = 0.005), hinting that CA3^CCK^ neurons are involved in the maintenance of hippocampal neuroplasticity.

### RNA interference of excitatory CA3^CCK^ expression impairs hippocampal functions

Next, to verify whether neuropeptide CCK in excitatory CA3^CCK^ neurons are also involved in spatial learning and memory, RNA interference (RNAi) strategy was utilized to downregulate CCK expression in CA3^CCK^ neurons (**Figure 6A**). We thus injected AAV9-CamkIIα-DIO-mCherry-shRNA (CCK) into area CA3 to knockdown target gene and AAV9-CamkIIα-DIO-mCherry-shRNA (Scramble) as control. The experimental group was Cre on anti-CCK to knock down the expression of CCK in the area CA3 of CCK Cre mice, while the control group was Cre on anti-scramble. We confirmed the expression of shRNAs in the hippocampus at 4 weeks after AAV injection (**Figure 6B**). Moreover, we further verified that anti-CCK shRNAs faithfully downregulated CCK mRNA levels *in vivo* by using the qPCR technique and histology method (**Figures 6C**; two sample t-test, df = 10, t = 4.95, anti-scramble 100 ± 7.03 % v.s. anti-CCK 58.73 ± 7.23 %, P = 0.001; **sFigure 4A-B**; two sample t-test, df = 16, t = 6.23, anti-scramble 52.44 ± 2.64 cells v.s. anti-CCK 19.12 ± 4.45 %, P < 0.001). Subsequently, we conducted the MWM task in the two groups of mice to assess the role of CA1-projecting CA3^CCK^ neurons in the hippocampus-dependent spatial learning (**Figure 6D**).

**Figure 6.**
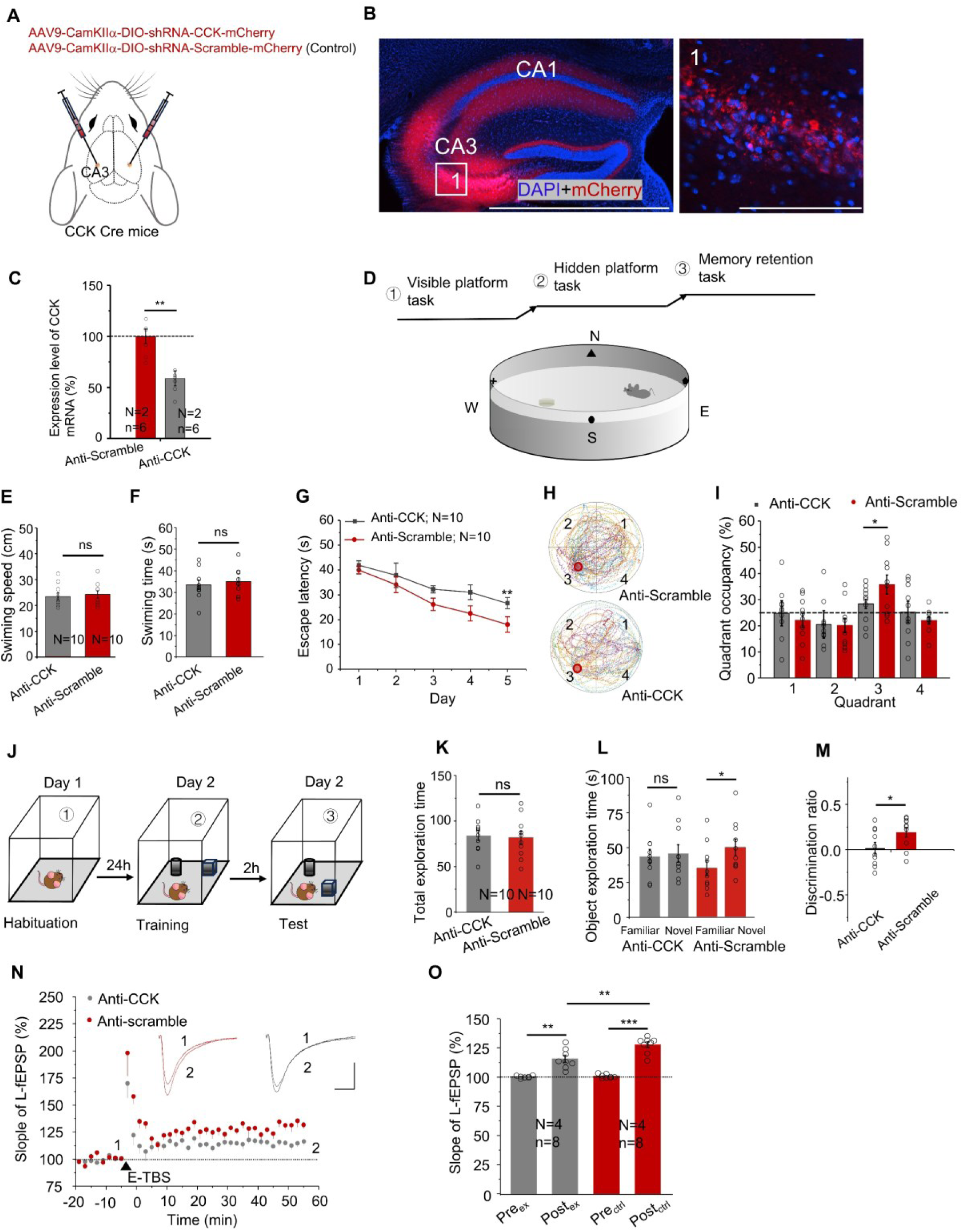
RNA interference of excitatory CA3^CCK^ expression attenuates hippocampal functions. Schematic illustrating the viral strategy for CCK knockdown in CA3 CCK-expressing neurons. CCK-Cre mice received injections of AAV9-CaMKIIα-DIO-(mCherry-bGH pA-U6)-shRNA-CCK or control AAV9-CaMKIIα-DIO-(mCherry-bGH pA-U6)-shRNA-Scramble (5.0 × 10¹² vg/mL, 300 nL) into the CA3 region. **(B)** Representative fluorescence images showing viral expression in the hippocampal CA3 region and labeled projections to the CA1 area. A higher-magnification image of the boxed region is shown on the right. Scale bars, 1000 µm (left) and 100 µm (right). **(C)** Quantification of CCK mRNA expression in the CA3 region of CCK-Cre mice infected with anti-CCK shRNA or scramble control shRNA, confirming efficient knockdown of CCK expression. **(D)** Schematic illustration of the hippocampal slice electrophysiological recording configuration. **(E)** Quantification of swimming speed during the visible platform task of the Morris water maze (MWM), showing no significant difference between anti-CCK and anti-scramble groups. **(F)** Quantification of latency to locate the visible platform, indicating intact sensorimotor performance in both groups. **(G)** Escape latency during the hidden platform training phase of the MWM for anti-CCK and anti-scramble groups. **(H)** Representative swimming trajectories of anti-CCK and anti-scramble mice during the spatial probe trial of the MWM. **(I)** Percentage of total time spent in each quadrant during the probe test, showing reduced preference for the target quadrant in anti-CCK mice compared with controls. **(J)** Schematic of the novel object location (NOL) task (see Methods for details). **(K)** Total object exploration time during the NOL task, showing no significant difference between anti-CCK and anti-scramble groups. **(L)** Control mice expressing scramble shRNA exhibited a greater preference for the object in the novel location compared with anti-CCK mice. **(M)** Quantification of discrimination index demonstrating significantly reduced spatial discrimination performance in anti-CCK mice relative to scramble controls. **(N)** Representative traces showing that long-term potentiation (LTP) induced by electrical stimulation was attenuated in hippocampal slices expressing anti-CCK shRNA compared with scramble controls. Scale bars, 0.2 mV and 10 ms. **(O)** Summary quantification of electrically evoked field excitatory postsynaptic potentials (E-fEPSPs) following LTP induction, demonstrating reduced synaptic potentiation in the anti-CCK group relative to controls. ∗p < 0.05, ^∗^∗p < 0.01, ∗∗∗p < 0.001; ns, not significant. Data are reported as mean ± SEM.

The two groups of mice injected with anti-CCK and anti-scramble showed comparable swimming speed (**Figure 6E**; two sample t-test, df = 18, t = −0.40, anti-CCK: 23.3 ± 2.75 cm/s v.s. anti-scramble: 24.1 ± 1.41cm/ s, p = 0.69) and swimming time in locating the visible platform above the water surface of the swimming pool (**Figure 6F**; two sample t-test, df = 18, t = −0.51, anti-CCK: 33.4 ± 2.21 s v.s. anti-scramble: 34.9 ± 1.93 s, p = 0.62). Then, mice infected with the anti-CCK showed deficits in spatial learning during the 5 days training (**Figure 6G**; two way mixed ANOVA, Bonferrori adjustment; F_4,15_ = 0.30, p = 0.88; anti-CCK 25.48 ± 2.12 s v.s. anti-scramble 18.05 ± 3.10 s on day 5, p = 0.007), and also exhibited deficiency in memory retention (**Figure 6H-I**; two way mixed ANOVA, Bonferrori adjustment; F_3,16_ = 1.20, p = 0.34; anti-CCK 28.38 ± 1.78 % v.s. anti-scramble 35.76 ± 3.68 % in quadrant 3, p = 0.03). These results suggest that knockdown of CCK expression in area CA3 attenuates spatial learning ability.

In another spatial learning protocol, the novel object location (NOL) task (**Figure 6J**), two groups of mice exhibited equal motivation to explore objects (**Figure 6K**; two sample t-test, df = 18, t = 0.28, anti-CCK: 83.82 ± 5.85 s v.s. anti-scramble: 81.22 ± 7.25 s, p = 0.78). Compared with the control mice (anti-scramble), the mice infected with anti-CCK failed to distinguish between the novel and familiar object location in the test phase, and displayed low discrimination ratio (**Figure 6L**; two-way mixed ANOVA, Bonferrori adjustment; F_1,18_ = 2.97, p = 0.10; anti-CCK: Familiar 42.63 ± 5.21 s v.s. Novel: 44.87 ± 6.33 s, p = 0.67; anti-scramble: Familiar 34.41 ± 5.30 s v.s. Novel: 49.22 ± 8.87 s, p = 0.01. **Figure 6M**; two sample t-test, df = 18, t = −2.0, anti-CCK: 0.02 ± 0.06 v.s. anti-scramble: 0.19 ± 0.05, p = 0.04), indicating that downregulation of CCK expression in area CA3 was sufficient to impair spatial memory. Taken together, these results support our hypothesis that excitatory CCK acts as a functional neuromodulator in the hippocampal system.

Additionally, for the electrophysiology recording, E-TBS adequately induced LTP at CA3-CA1 synapses in control slices containing the anti-scramble shRNAs, but the same protocol elicited remarkably smaller LTP on slices with anti-CCK shRNA targeting the sequences of CCK gene (**Figures 6N-O**; two way mixed ANOVA, Bonferrori adjustment; F_1,14_ = 10.06, p = 0.007; Pre_ctrl_: 100.93 ± 0.57 % v.s. Post_ctrl_: 127.78 ± 2.47 %, p < 0.001; Pre_ex_: 100.44 ± 0.34 % v.s. Post_ex_: 115.76 ± 2.96 %, p < 0.001; Post_ctrl_ v.s. Post_ex,_ p = 0.008). Moreover, we noticed that the amplitude of the electrically evoked fEPSPs in anti-CCK group is similar to those of the control group (**Figures 6N**, insert), suggesting normal neurotransmission in hippocampal slices from both groups. Therefore, the impairment of LTP is likely due to the reduced expression of CCK in CA1-projecting CA3^CCK^ terminals which impaired the neuromodulation.

## Discussion

In the present study, we reported the distribution profile of CCK positive neurons in the dorsal hippocampus, a key brain region associated with spatial memory formation and transformation (Martin and Clark 2007). Although the role of inhibitory CCK neurons is well studied in the hippocampus, the function of excitatory CCK neurons is still unclear. Optogenetic activation of the CCK inhibitory populations produces a prominent IPSC and controls the input and output gain of CA1 pyramidal neurons, and this kind of modulation is mediated by the presynaptic CB1 receptors (Hartzell et al., 2018). A recent study uncovered that systemic activation of CCK-GABA neurons minimally affects emotion but significantly enhances cognition and memory (Whissell et al., 2019). In our study, we combined the Ai14::CCK Cre mice and immunochemistry method to show the distribution of CCK positive neurons in the hippocampus. Moreover, we adopted the pharmacological approach to diminish the tendency for light-evoked fEPSPs of excitatory CA1-projecting CA3^CCK^neurons. Additionally, using the patch clamp technique to record the EPSP of these neurons also can directly validate this conclusion. Due to technical limitations at the current stage, we were unable to perform whole-cell recordings or pharmacological manipulations using CCK receptor antagonists. In future studies, the application of these approaches to directly record and selectively block EPSPs from excitatory CCK neurons in the hippocampus will further strengthen and validate our conclusions. Importantly, to further improve cell-type specificity, we propose an intersectional genetic strategy using CCK-IRES-Cre × VGlut1-Flp mice combined with a Cre-On/Flp-On (Con/Fon) AAV, which would restrict expression exclusively to excitatory CCK-expressing neurons and eliminate potential contributions from inhibitory CCKL cells. This approach will be implemented in future studies to refine circuit specificity. To our knowledge, this is the first study that demonstrated the function of excitatory CCK neurons in the hippocampus.

Additionally, to fully validate the involvement of excitatory CA1-projecting CA3^CCK^ neurons in hippocampal processing involving learning and neuroplasticity, we used the chemogenetic technique to examine the function of CA3^CCK^ neurons. Interestingly, we found that specific inhibition of these neurons significantly impaired the CA1-CA3 LTP and attenuated spatial learning. Many studies have also demonstrated that neurotransmitters facilitate the hippocampal plasticity and further modulate the behavior performance (Birthelmer et al., 2003; Ohta et al., 2003; Lopes et al., 2002), because deficiency of neurotransmitters in the CNS is known to be able to decrease the extent of synaptic transmission in the CA3-CA1 pathway. For instance, Mlinar reported that endogenous 5-hydroxytryptamine (5-HT) potentiates the LTP in the hippocampus and its positive effects on cognitive performance (Mlinar et al., 2015). Also, another study used a viral-mediated approach to delete brain-derived neurotrophic factor (BDNF) specifically in CA3-CA1 pathway, and further demonstrated that presynaptic and postsynaptic BNDF are essential for LTP induction and maintenance, as well as contextual memory impairments (Mohajerani et al., 2007). In this study, we used RNA interference technique (RNAi) to fully characterize the function of CA3^CCK^ neurons. Intriguingly, the knock down of CCK expression significantly impaired LTP formation at excitatory CA1-projecting CA3^CCK^ synapses and further affected the hippocampus dependent-spatial learning and memory. Interestingly, an earlier study demonstrated that intraperitoneal injection of exogenous CCK-4 significantly improved performance in hippocampus-dependent spatial learning tasks in both CCK gene knockout (CCK-KO) mice and Alzheimer’s disease (AD) mouse models (Zhang et al., 2024). These findings suggest that enhancing CCK signaling can ameliorate hippocampal dysfunction at both the behavioral and synaptic plasticity levels. Therefore, we established the essential role of CCK as an active neuromodulator that regulates hippocampal plasticity and its dependent behaviors (**Figure 7**).

**Figure 7.**
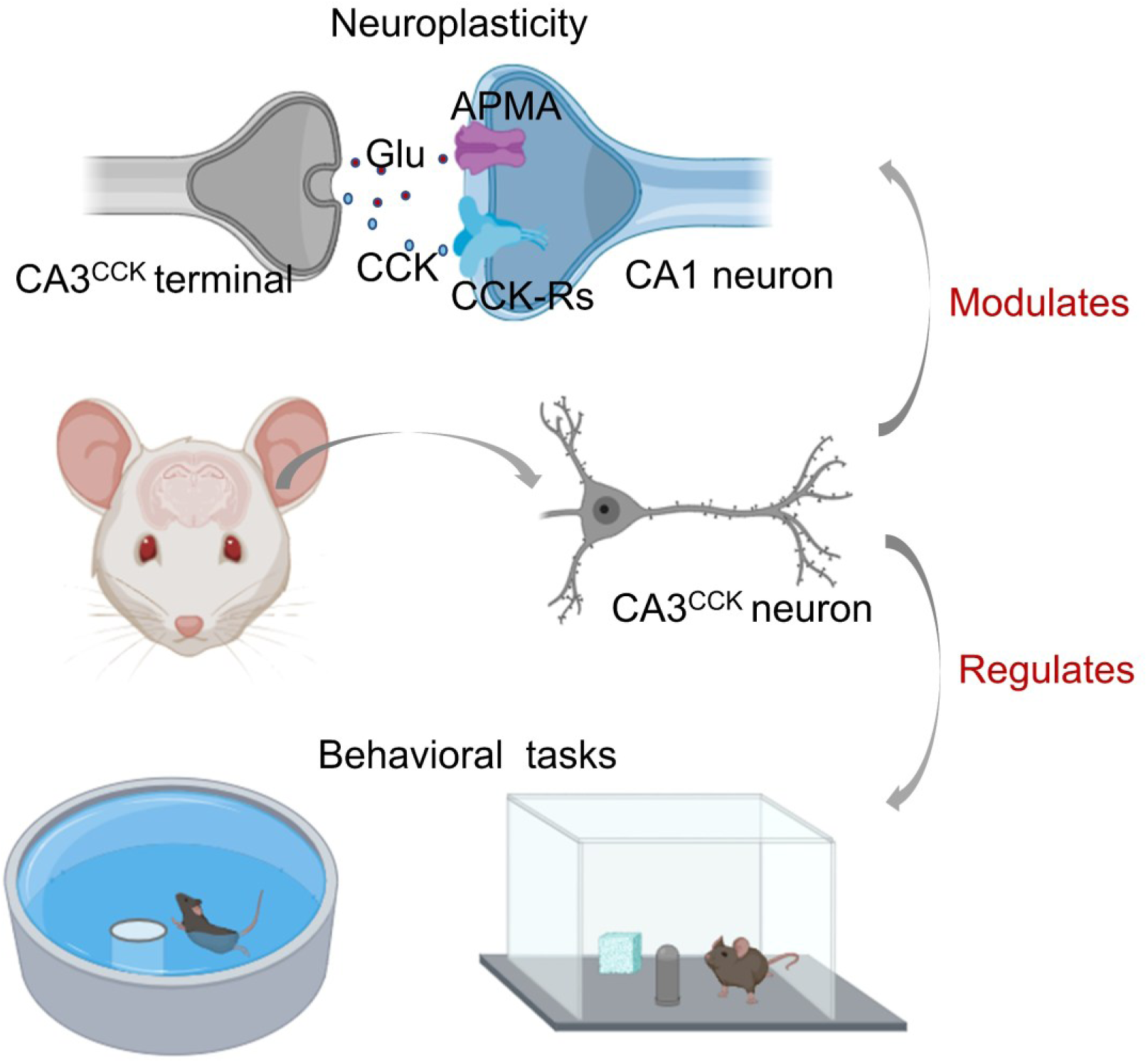
Summary model illustrating the role of the perforant pathway in regulating spatial memory and hippocampal synaptic plasticity

Establishing a causal link between neural plasticity and spatial learning is crucial for understanding the status of CA3^CCK^ neurons in the hippocampus. Our investigation verified that CA3^CCK^ neurons modulate hippocampal long-term plasticity in a homosynaptic manner, and their activity underlies spatial learning and memory formation. We demonstrated that the CA3^CCK^ projections to the CA1 region are required and sufficient to induce the LTP at CA3-CA1 synapses. In the behavioral setup, we also observed that the calcium response of CA1-projecting CA3^CCK^ neurons increased during the exploration. The Ca^2+^ activities were also raised throughout the MWM task. Nevertheless, perturbation of the activities of CA3^CCK^ neurons impaired spatial learning both in the NOL task and MWM task. It was likely that the CA3 input to the CA1 conveys spatial information. Thus, we can conclude that CA3^CCK^ neurons may specifically regulate the hippocampal state and plasticity during behavioral performance.

Finally, further experimental works could be undertaken to investigate the function of CA3^CCK^ neurons. For instance, the change of CCK concentration during the light high frequency stimulation of CA1-projecting CA3^CCK^ neurons by using the GPCR-based CCK sensor. Several high-sensitive GPCR-based sensors for detecting the neurotransmitters have been well developed, including the dopamine, norepinephrine, serotonin and acetylcholine. Besides, using a GPCR-based CCK-BR sensor combined with fiber photometry, our previous work demonstrated rapid, activity-dependent CCK release in the hippocampus during object-exploratory behavior, supporting a functional role for hippocampal CCK signaling in cognitive tasks (Su et al., 2023). Given that hippocampal neurons receive CCK-positive projections from multiple brain regions, it remains technically challenging to precisely identify the cellular source of CCK release in CA1 during behavior. Future studies employing selective CCK overexpression in CA3 neurons, together with CCK-BR sensor recordings, may help further delineate the contribution of CA3-derived CCK to hippocampal-dependent behaviors.

## Methods

### Animals

Adult Cck-IRES-Cre (CCK Cre: RRID:IMSR_JAX:012706) mice and Ai14 mice (RRID:IMSR_JAX:007914) were used in this study. In behavioral experiments, all male mice were housed in a 12-h light/dark cycle and were provided food and water ad libitum. All experimental procedures were approved by the Animal Subjects Ethics Sub-Committee of the City University of Hong Kong.

### Viruses

Adeno-associated virus (AAVs) were purchased from the BrainVTA, Wuhan, China: AAV9-CaMKIIa-DIO-GCaMP6S-WPRE-hGH-pA (PT-0090); AAV9-CaMKIIa-DIO-(mCherry-bGH pA-U6)-shRNA(CCK)-WPRE-hGH pA (PT-9086); AAV9-CaMKIIa-DIO-(mCherry-bGH pA-U6)-shRNA(Scramble)-WPRE-hGH pA (PT-3088); AAV9-CaMKIIa-DIO-hM4D(Gi)-mCherry-WPRE-hGH polyA (PT-1143); AAV9-Syn-DIO-mCherry-WPRE-hGH polyA (PT-0115); AAV9-hSyn-CCK sensor 2.3-GFP; AAV9-hSyn-GFP; Addgene, Cambridge, MA, USA: Retrograde AAV-EF1a-DIO-EYFP (27056). Taitool BioScience, Shanghai, China: AAV9-mCaMKIIa-DIO-ChrimsonR-mCherry-ER2-WPRE-pA (S0728-9).

### AAV injection

Adult mice were deeply anesthetized with an intraperitoneal injection of pentobarbital sodium (50 mg/kg; Ceva Santé Animale, France) and secured in a stereotaxic frame. Following a midline scalp incision to expose the skull, the stereotaxic apparatus was adjusted to ensure that bregma and lambda were positioned in the same horizontal plane. A small craniotomy was drilled above the target site, and adeno-associated virus (AAV) was unilaterally delivered into the CA3 region of the hippocampus using a nanoliter injector (Micro4 system, World Precision Instruments) fitted with a quartz glass micropipette. The stereotaxic coordinates relative to bregma were as follows: anteroposterior (AP), −1.70 mm; mediolateral (ML), +2.35 mm; dorsoventral (DV), −1.85 mm.

The viral suspension was infused at a constant rate of 30 nL/min. The total injection volume and viral titer are specified in the corresponding figure legends. To minimize backflow and ensure proper diffusion of the virus, the micropipette was left in place for 5 minutes after completion of the infusion before being slowly withdrawn. After injection, the incision was sutured and treated with erythromycin ointment to prevent infection. Mice were placed on a heating pad until fully recovered from anesthesia and then returned to their home cages for postoperative care.

### Brian slice recording

For acute brain slice preparation, mice were deeply anesthetized with 1.4% gaseous isoflurane (Wellona Pharma) and decapitated. The brain was rapidly removed and immediately immersed in ice-cold, oxygenated artificial cerebrospinal fluid (aCSF; 95% OL / 5% CO□). The aCSF contained (in mM): 124 NaCl, 3 KCl, 1.25 KH□PO□, 1.25 MgSO□, 2 CaC□□, 26 NaHCO□, and 10 glucose, with a pH of approximately 7.4. This same aCSF solution was used throughout all subsequent procedures, including tissue dissection, slicing, incubation, and electrophysiological recording. Coronal brain slices (300 µm thickness) containing the hippocampus were prepared using a vibrating microtome (Leica VT1000S). Slices were transferred to oxygenated aCSF and allowed to recover at 32°C before recording.

For electrophysiological recordings, individual hippocampal slices were placed onto a multi-electrode array (MEA) probe and continuously perfused with oxygenated aCSF. Field excitatory postsynaptic potentials (fEPSPs) were recorded using an MEA recording system (Alpha MED Sciences) integrated with a light stimulation module (Inper), allowing for both electrically and optically evoked responses. For electrical stimulation, a designated electrode on the MEA probe served as the stimulating electrode. For optical stimulation, a 2-ms pulse of red light (635 nm) was delivered through a 200-µm-diameter optical fiber positioned over the CA1 region. After stable baseline fEPSP responses were established, input–output (I/O) curves were generated by plotting the slope of the fEPSP against increasing electrical stimulus intensities or optical light intensities. fEPSP slopes were normalized to baseline responses and analyzed using MED Mobius software (Alpha MED Sciences).

### Real-time PCR

Tissue samples from the target brain region (hippocampal CA3) were freshly dissected from mice four weeks after AAV-mediated expression of anti-CCK shRNA or scramble control shRNA. Bilateral CA3 tissue was collected for RNA analysis. Total RNA was extracted using TRIzol reagent according to the manufacturer’s instructions. Extracted RNA was reverse transcribed into complementary DNA (cDNA) using Maxima Reverse Transcriptase (Thermo Fisher Scientific). Quantitative real-time PCR was performed using a QuantStudio™ 3 Real-Time PCR System (Applied Biosystems) to specifically amplify the CCK transcript. The primer sequences used for CCK amplification were as follows: CCK forward, 5′-AGCGCGATACATCCAGCAG-3′; CCK reverse, 5′-ACGATGGGTATTCGTAGTCCTC-3′. Relative CCK mRNA expression levels were quantified using the comparative threshold cycle (Ct) method (2^−ΔΔCt). Expression levels were normalized to the housekeeping gene GAPDH. Three biological replicates were included for each experimental group.

### Morris water maze task

The Morris water maze (MWM) task was conducted according to the protocol described by Vorhees and Williams (2006). The apparatus consisted of a circular pool with a diameter of 122 cm filled with opaque water. All behavioral testing was performed under consistent lighting conditions, with illumination set sufficiently bright to allow accurate video tracking. Prior to hidden platform training, all mice were subjected to a visible platform task to assess visual acuity and swimming ability. During this phase, mice were allowed to locate a visible platform placed above the water surface.

For spatial learning, mice underwent hidden platform training for five consecutive days. During each training session, the platform was submerged below the water surface and remained in a fixed location. Mice were released into the pool and allowed up to 60 s to locate the hidden platform. If a mouse failed to find the platform within the allotted time, it was gently guided to the platform. Mice were allowed to remain on the platform briefly before being returned to their home cages. On day 6, a probe trial was performed to assess spatial reference memory. The platform was removed from the pool, and mice were allowed to swim freely to evaluate memory retention. The former platform location was in the target quadrant (quadrant 3). Swimming trajectories, time spent in each quadrant, and search patterns were recorded and analyzed using MATLAB-based tracking software.

### Novel object location task

The novel object location (NOL) task was conducted as previously described (Pérez-García et al., 2016) and consisted of three sequential phases. On day 1, each mouse was habituated to the empty testing apparatus for 10 min. On day 2, mice underwent a training phase followed by a testing phase. During the training phase, mice were placed in the apparatus containing two identical objects positioned in fixed locations and were allowed to freely explore for 5 min. After training, mice were returned to their home cages for a 1-h retention interval.

For the testing phase, mice were reintroduced into the same apparatus for 5 min. One of the two objects remained in its original location (familiar object location, FOL), whereas the other object was moved to a novel location within the arena (novel object location, NOL). Object identity remained unchanged between training and testing. Exploration behavior was recorded, and exploration time for each object was quantified. The discrimination index (DI) was calculated using the following formula: (Pérez-García et al., 2016):

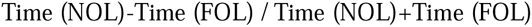

A positive DI value indicates a preference for the object in the novel location, a negative value indicates a preference for the familiar location, and a value of zero indicates equal exploration of both locations.

### Chemogenetic manipulation

To enable chemogenetic manipulation of CCK-expressing neurons, CCK-Cre mice were bilaterally implanted with guide cannulas targeting the CA1 region of the hippocampus following viral injection. Stereotaxic coordinates relative to bregma were as follows: anteroposterior (AP), −1.75 mm; mediolateral (ML), ± 1.25 mm; dorsoventral (DV), −1.20 mm. The surgical procedures for cannula implantation were identical to those described above.

Four weeks after viral injection, mice underwent hippocampal-dependent behavioral testing, including the Morris water maze (MWM) and novel object location (NOL) tasks, as well as electrophysiological recordings. Prior to behavioral testing, mice expressing the inhibitory DREADD hM4D(Gi) and control mice expressing mCherry received bilateral microinfusions of clozapine-N-oxide (CNO; 10 μM in 0.1% DMSO) into the CA1 region. For each infusion, 300 nL of CNO solution was delivered per side at a rate of 50 nL/min using a programmable microinjector.

For in vitro electrophysiological experiments, CNO (50 μM) was bath-applied by adding it to the perfusing artificial cerebrospinal fluid (aCSF) during field excitatory postsynaptic potential (fEPSP) recording sessions.

### Fiber photometry recording

Fiber photometry recordings were performed using a commercially available system (Doric Lenses). Sinusoidally modulated light-emitting diodes (LEDs) at 473 nm (220 Hz) and 405 nm (330 Hz) were used to excite the calcium-dependent GCaMP fluorescence signal and the calcium-independent isosbestic control signal, respectively. Excitation light was delivered through an implanted optical fiber, with the output power calibrated to 10 μW at the fiber tip. Fluorescence emission was collected through the same optical fiber and directed to two independent photoreceivers (Model 2151, Newport Corporation). LED modulation, signal acquisition, and real-time demodulation of the 473 nm and 405 nm fluorescence signals were controlled by an RZ5P data acquisition system equipped with a real-time processor (Tucker-Davis Technologies). During behavioral experiments, object exploration in the novel object location (NOL) task was defined as the mouse approaching, sniffing, or touching an object. To enable concurrent fiber photometry recordings, a simplified version of the Morris water maze (MWM) task was used, as previously described (Qin et al., 2018). All photometry data were analyzed using the open-source pMAT software package (Bruno et al., 2021). The raw 473 nm GCaMP signal was corrected for motion-related artifacts by fitting the 405 nm isosbestic signal to the 473 nm channel. The normalized calcium signal was expressed as ΔF/F and calculated using the following formula: ΔF/F = (473 nm signal - fitted 405 nm signal) / fitted 405 nm signal.

### Anatomy and histology

All animals were anesthetized with sodium pentobarbital and were perfused with phosphate-buffered saline and fixed with paraformaldehyde solution. Then, mouse’s brain was detached and submerged into 4 % PFA solution for fixations at 4 °C in the refrigerator. The brains were sectioned into 40 mm-thick slices via the vibratome machine. The prepared brain sections were counter-stained with DAPI (1:10000) and mounted onto grass slides with 70% glycerol in PBS to observe the AAV expression and fiber track. Finally, the coverslips were used to cover the brain slices and sealed with adhesive glue.

For the histological procedure, brain slices were washed with 0.01 M PBS and blocked with blocking solution (15 % goat serum mixed with 0.4 % Triton X-100) at room temperature for 2-3 h. Then, the primary antibody was prepared and incubated with brain slices in 24-well cell culture plates overnight at 4 °C. Then, brain slices were washed by 0.01 M PBS and coupled to a secondary antibody at room temperature for 2-3 h. Next, brain slices were washed with PBS and were stained with DAPI. Finally, fluorescence image of brain slices was captured by using a Nikon Eclipse fluorescence microscope and a Nikon A1HD25 confocal microscope.

### Statistical analysis

All statistical analyses (including two sample t-test, and two-way mixed ANOVA) were done in SPSS (IBM, USA). Statistical significance was set at p < 0.05.

## Author contributions

Conceptualization: H.F; Methodology: H.F; Investigation: H.F and A.B; Writing: H.F. and S.T.B.

## Data availability statement

The data that support the findings of this study are available from the corresponding author upon reasonable request

## Acknowledgements

We thank Prof. Jufang He for providing resources. This work was supported by funding from the following: Hong Kong Research Grants Council, General Research Fund: CityUHK 11101521, CityUHK 11103922, CityUHK 11104923, CityUHK 11104524.

## Supplementary Figures

**sFigure1.**
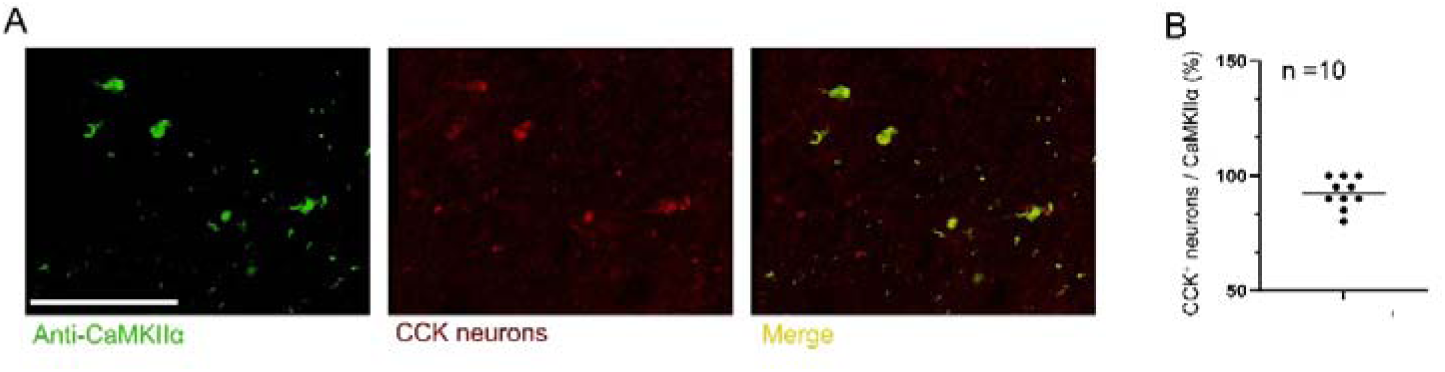
Excitatory CA3 neurons secret the neuropeptide CCK. **(A)** Representative fluorescent images show that colocalization of the CaMKIIα and CCK^+^ neurons. Scale bar = 50 µm. **(B)** Quantitative analysis shows the ratio between CCK^+^ neurons and CaMKIIα in the CA3 area.

**sFigure2.**
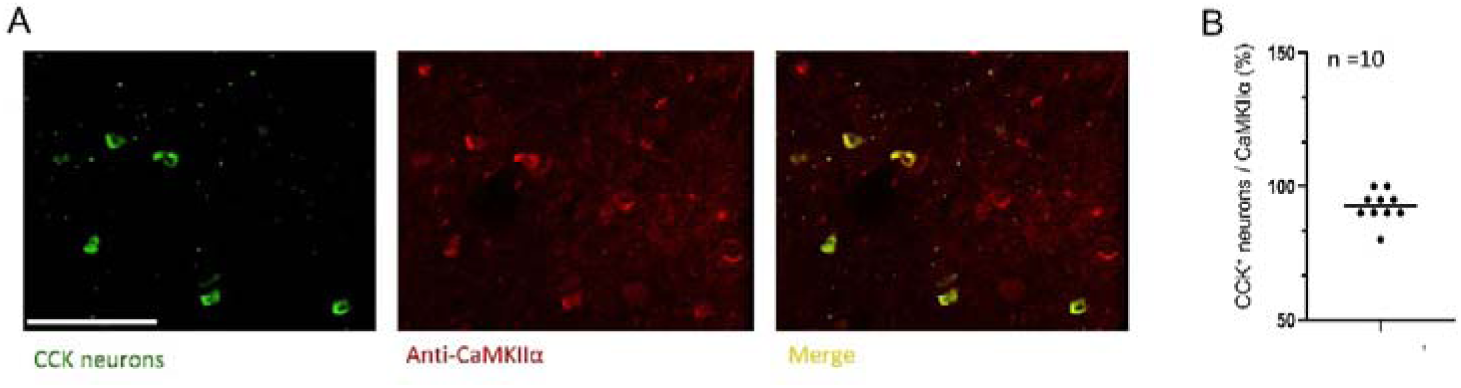
CA3^CCK^ neurons fire actively during hippocampal-dependent tasks. **(A)** Representative fluorescence images showing colocalization of CCK-positive neurons with the excitatory neuronal marker CaMKIIα in the CA3 region. Scale bar = 50 µm. **(B)** Quantification of the proportion of CCK-positive neurons that co-express CaMKIIα in the CA3 area.

**sFigure3.**
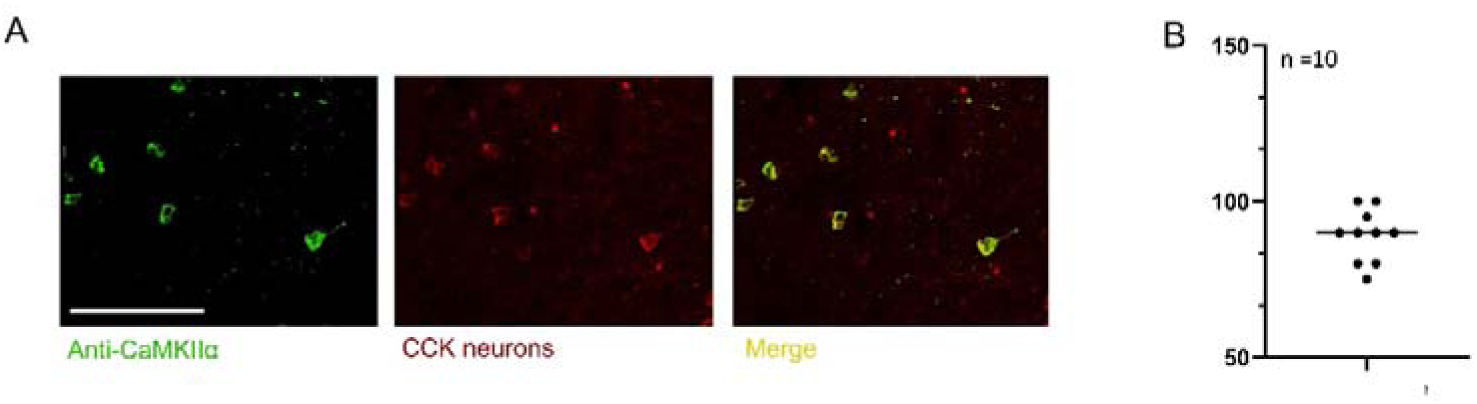
Chemogenetic inhibition of the excitatory CA3^CCK^-CA1 pathway impairs behavioral tasks. **(A)** Representative fluorescent images show that colocalization of the CaMKIIα and CCK neurons. Scale bar = 50 µm. **(B)** Quantitative analysis shows the ratio between CCK neurons and CaMKIIα in the CA3 area.

**sFigure4.**
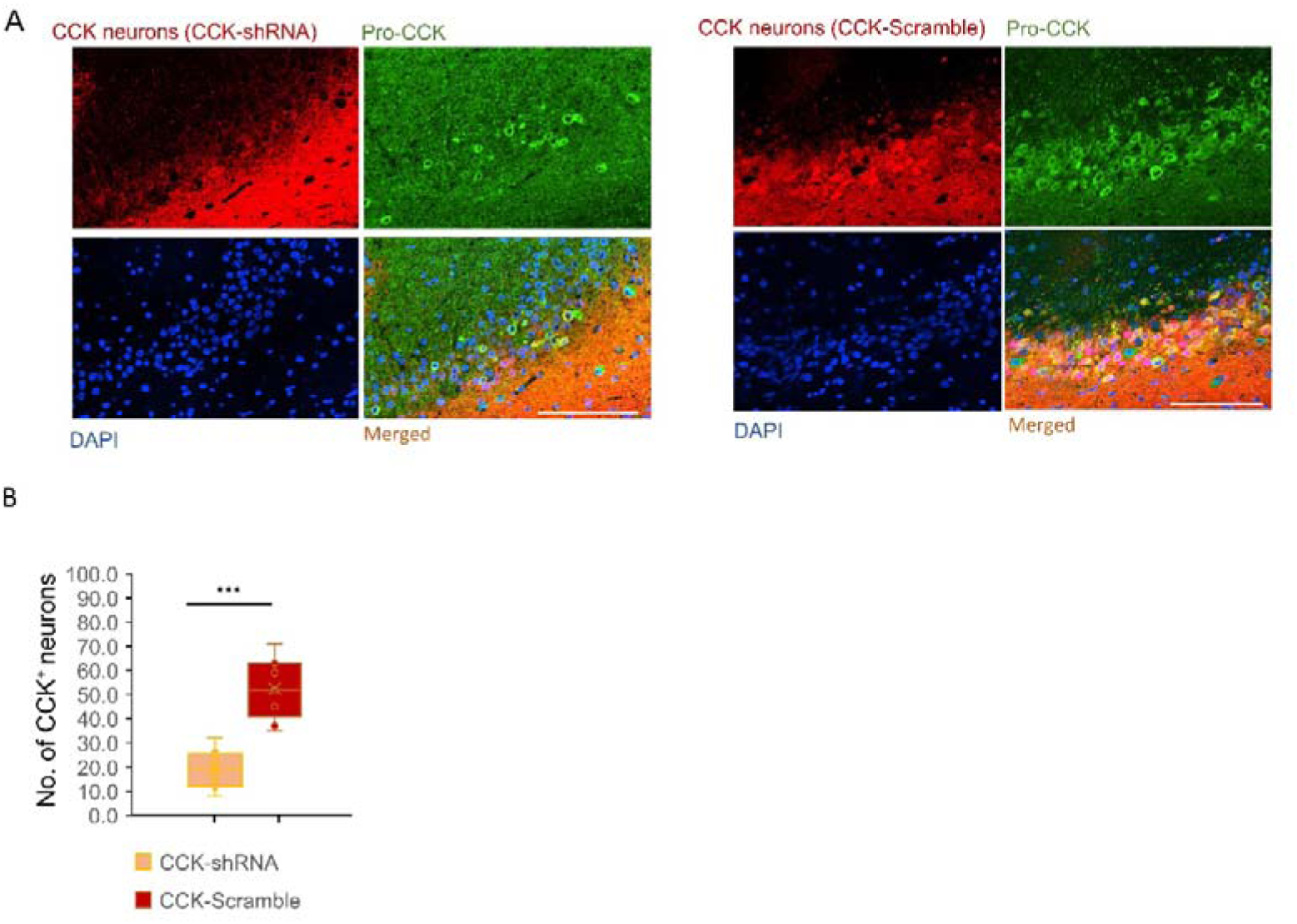
RNA interference of excitatory CA3^CCK^expression attenuates hippocampal functions. **(A)** Representative fluorescent images show that CCK-shRNA (left panel) significantly reduced CCK expression in CA3 -positive neurons compared with the CCK-Scramble group (right panel). Scale bar = 50 µm. **(B)** Quantitative analysis shows the number of CCK neurons, defined by colocalization of CCK and Pro-CCK, in the CCK-shRNA and CCK-Scramble groups (n = 9 slices for each group). ∗p < 0.05, ∗∗p < 0.01, ∗∗∗p < 0.001; ns, not significant. Data are reported as mean ± SEM.

