## Supplementaty Figures for "Excitatory cholecystokinin neurons in CA3 area regulate the navigation learning and neuroplasticity"

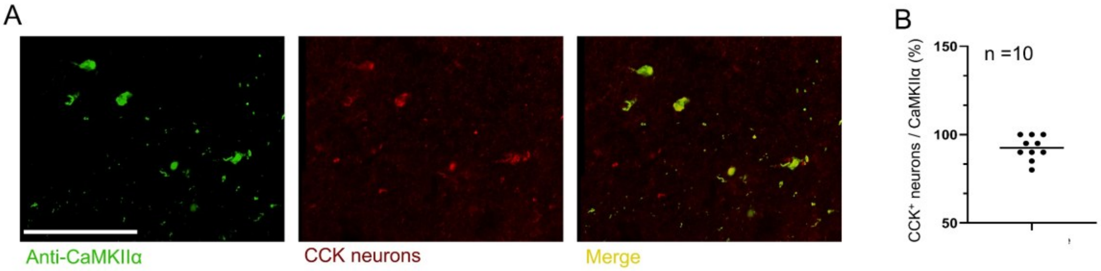


**sFigure1. Excitatory CA3 neurons secret the neuropeptide CCK.**

A. Representative fluorescent images show that colocalization of the CaMKIIα and $CCK^+$ neurons. Scale bar = 50 μm.
B. Quantitative analysis shows the ratio between $CCK^+$ neurons and CaMKIIα in the CA3 area.

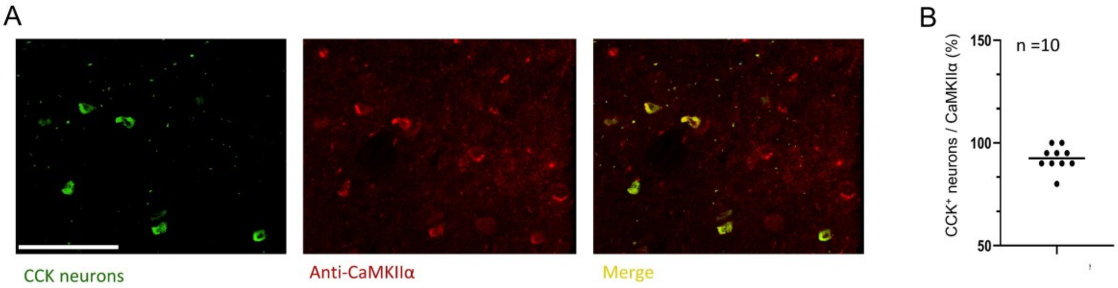


**sFigure2. CA3$^{CCK}$ neurons fire actively during hippocampal-dependent tasks.**

A. Representative fluorescence images showing colocalization of CCK-positive neurons with the excitatory neuronal marker CaMKIIα in the CA3 region. Scale bar = 50 μm.
B. Quantification of the proportion of CCK-positive neurons that co-express CaMKIIα in the CA3 area.

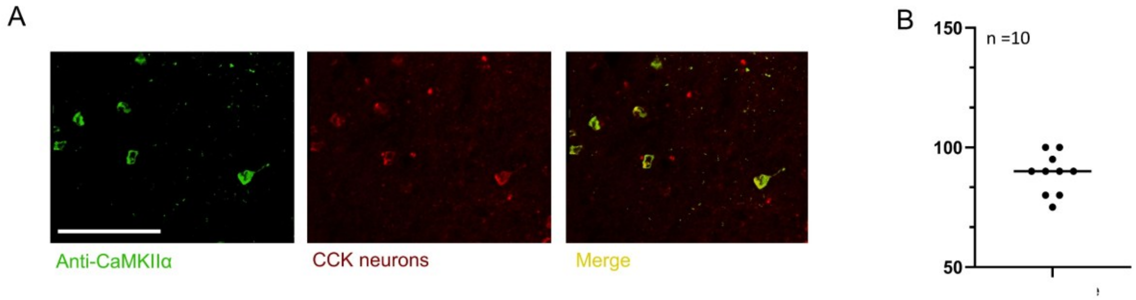


**sFigure3. Chemogenetic inhibition of the excitatory CA3$^{CCK}$-CA1 pathway impairs behavioral tasks.**

A. Representative fluorescent images show that colocalization of the CaMKIIα and CCK$^{+}$ neurons. Scale bar = 50 μm.
B. Quantitative analysis shows the ratio between CCK$^{+}$ neurons and CaMKIIα in the CA3 area.

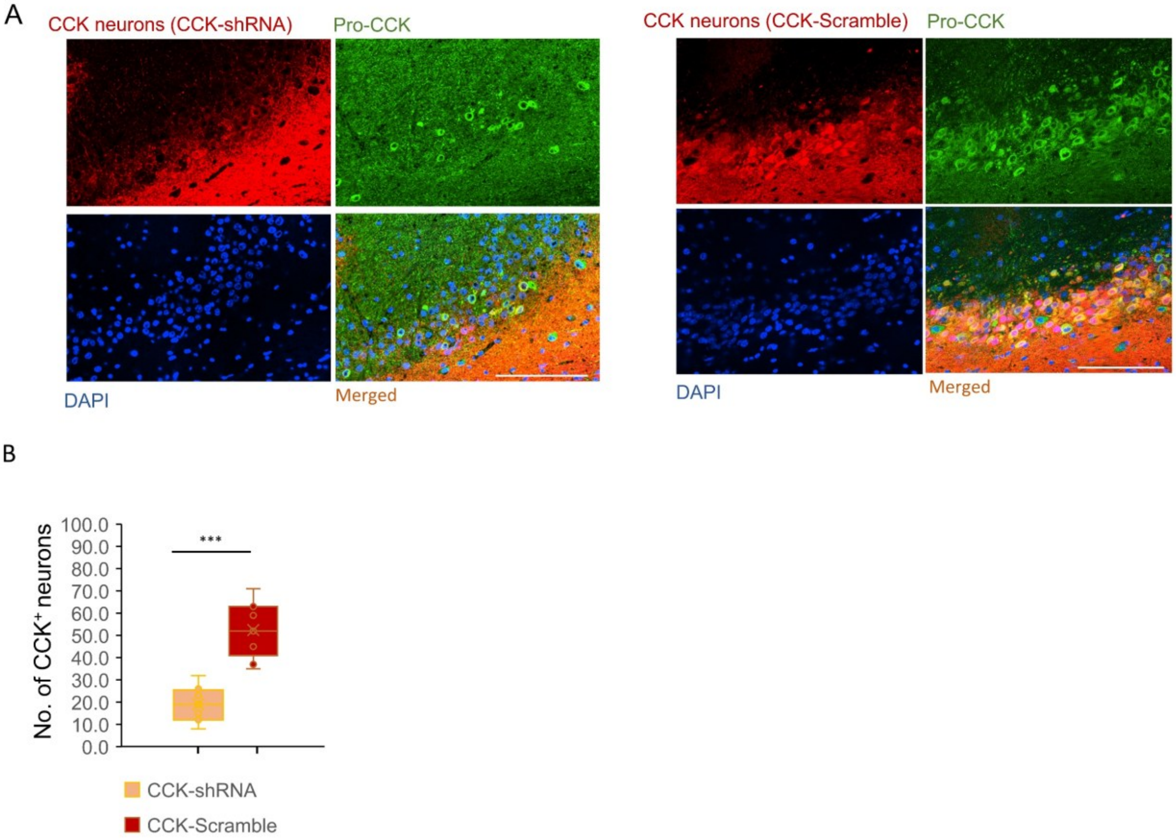


**sFigure4. RNA interference of excitatory $CA3^{CCK}$ expression attenuates hippocampal functions.**

A. Representative fluorescent images show that CCK-shRNA (left panel) significantly reduced CCK expression in $CA3^{CCK}$-positive neurons compared with the CCK-Scramble group (right panel). Scale bar = 50 μm.

B. Quantitative analysis shows the number of CCK neurons, defined by colocalization of CCK and Pro-CCK, in the CCK-shRNA and CCK-Scramble groups (n = 9 slices for each group).

$*p < 0.05$, $**p < 0.01$, $***p < 0.001$; ns, not significant. Data are reported as mean ± SEM.
